# CADGE 2.0, Transcription-Translation-Coupled DNA Replication is Improved in a Chemically Modified Cell-Free System

**DOI:** 10.64898/2026.02.27.708527

**Authors:** Riku Nagai, Carlos Chavez Ramirez, Zhanar Abil

## Abstract

*In vitro* directed evolution in synthetic microcompartments can generally support the evolution of genes with functions beyond affinity. The main challenge in the implementation of this strategy is the need to incorporate no more than a single DNA template molecule per microcompartment, thereby establishing a robust genotype-phenotype linkage, but which results in slow, inconsistent *in vitro* transcription and translation (IVTT) and poor DNA recovery after selection or screening. To address this challenge, we previously developed CADGE (Clonal Amplification-enhanceD Gene Expression) a strategy that allows the clonal amplification of linear gene-encoding DNA and coupled, *in situ* transcription-translation of the gene of interest. Here, we show that clonal amplification is highly sensitive to the cell-free system’s composition and that robust, highly efficient cell-free DNA amplification via the CADGE platform can be achieved by replacing standard vendor-supplied energy mixes with DNA replication-optimized, homemade counterparts.

## Introduction

Directed evolution is a powerful tool in synthetic biology.^1–3^ Yet, despite numerous achievements, it remains largely limited to cell-based systems. Cell-free directed evolution offers an alternative that enables the evolution of toxic or growth-incompatible functions and allows selection to act directly on molecular activity rather than organismal fitness.^4–7^ Such approaches rely on cell-free gene expression systems that can support *in vitro* transcription-translation (IVTT) of proteins^8^ in an open, chemically tunable environment.^9^ A common strategy to establish genotype–phenotype linkage in cell-free evolution is *in vitro* compartmentalization.^6,10,11^ Genotype–phenotype linkage can be achieved by using low initial DNA copy number so that each compartment contains, at most, a single variant.^11^ However, low template concentrations often result in slow and variable IVTT,^12–16^ and DNA recovery from microcompartments remains a major bottleneck in both *in vitro* and cell-based systems.^17–19^ To overcome these limitations, various strategies for clonal DNA amplification in microcompartments have been proposed, but these approaches require cumbersome procedures or specialized instrumentation, such as bead display or microfluidic droplet generation.^20–27^

In a previous study, we developed CADGE (Clonal Amplification-enhanceD Gene Expression), a highly streamlined alternative to existing clonal DNA amplification strategies with the potential to accelerate *in vitro* evolution.^28^ This strategy combines, within a single compatible environment, isothermal amplification of a template DNA starting from a single copy with *in vitro* transcription-translation, followed by selection or screening.^28^ DNA amplification is carried out by the minimal DNA replication machinery from the *B. subtilis* phage Φ29.^29^ The machinery consists of DNA polymerase (DNAP; *p2* gene), terminal protein (TP; *p3* gene), double-stranded DNA-binding protein (DSB), and single-stranded DNA-binding protein (SSB). Replication initiates independently at two origins (*ori*)^30,31^ located at the termini of the linear Φ29 genome: DSB binds to *ori*^32,33^ and recruits the DNAP–TP complex,^34^ TP primes synthesis,^35^ and DNAP simultaneously replicates^36^ and displaces the non-template strand,^37^ which is stabilized by SSB.^38^ Two DNAP molecules thus replicate both strands from opposite ends.^39^ CADGE repurposes this protein-primed system by placing the gene of interest between two *ori* sites on a linear template, with DSB and SSB supplied recombinantly and DNAP and TP provided either as purified proteins or expressed *in situ*.^28^ Unlike rolling-circle amplification,^40–43^ protein-primed replication^44,45^ regenerates linear DNA identical to the template, minimizing DNA handling between evolution rounds. When encapsulated in liposomes, CADGE boosts phenotypic output under low-copy conditions and improves DNA recovery after selection or screening.^28^

CADGE was established in the recombinant cell-free system, PURE (Protein synthesis Using Recombinant Elements).^46^ However, we and others^47^ recently observed that commercially available PURE systems currently demonstrate significantly decreased levels of DNA replication compared to previously reported levels,^28^ revealing strong sensitivity to modifications in PURE composition. We hypothesized that proprietary changes in reaction composition that optimize transcription and translation can tip the balance against replication. Several studies successfully re-engineered PURE formulations to balance IVTT with Φ29 DNAP–dependent DNA replication from circular templates, particularly by adjusting NTP and tRNA concentrations and rebalancing Mg^2+^ to account for NTP-dependent chelation.^42,43,48,49^ However, it was unclear whether formulations optimized for nucleic-acid-primed rolling-circle amplification would also support protein-primed replication of linear templates, which requires TP–dAMP initiation and a distinct transition from initiation to elongation.^50,51^ These mechanistic and kinetic differences make direct extrapolation nontrivial. Consistent with this concern, despite prior demonstrations of robust Φ29-mediated rolling-circle replication in extract-based systems,^49^ we were unable to implement CADGE in crude *E. coli* lysate (data not shown). Together, these observations motivated us to test whether previously reported PURE formulations that support rolling-circle amplification could be adapted to balance IVTT with protein-primed Φ29 replication in CADGE.

Here, we demonstrate that PURE energy mix (the small-molecule components of the PURE kit) previously optimized^42^ for transcription and translation-coupled DNA replication markedly improves protein-primed Φ29 DNA replication from linear templates across commercial PURE platforms, while maintaining transcription–translation activity. We also observed a modest improvement in autocatalytic self-replication of the Φ29 DNA polymerase gene.^45,52^ This gene circuit can be integrated with diverse genetic modules to enable systems-level evolution and advance the construction of synthetic cells.^53^ Together, these results provide a practical foundation for protein-primed *in vitro* transcription-translation-DNA replication (IVTTR) using linear DNA that lower technical barriers to cell-free directed evolution of proteins and advance efforts toward building evolvable synthetic cells.

## Results and Discussion

### DNA-replication-optimized energy mix restores CADGE activity in PUREfrex

To restore CADGE in commercial PURE systems, we tested whether the DNA replication-optimized energy mix^42^ improves DNA replication in CADGE (Fig. 1A, 1B). For easy readout of DNA amplification and coupled transcription and translation, we used the yellow fluorescent protein (YFP) as a reporter. We used the *yfp* gene encoded on a linear DNA template and flanked by *ori* sequences at both ends (*ori-yfp*). The *yfp* DNA template was introduced in the CADGE reaction at a final concentration of 10 pM. Φ29 DNAP and TP were expressed by IVTT *in situ* from a plasmid DNA template. We expected that successful replication would result in increased *yfp* gene concentration, which would in turn elevate the rate of IVTT and yield of the YFP protein. YFP expression and DNA replication were monitored across three commercially available PURE platforms: PURExpress, PUREfrex 1.0, and PUREfrex 2.0.

**Figure 1.**
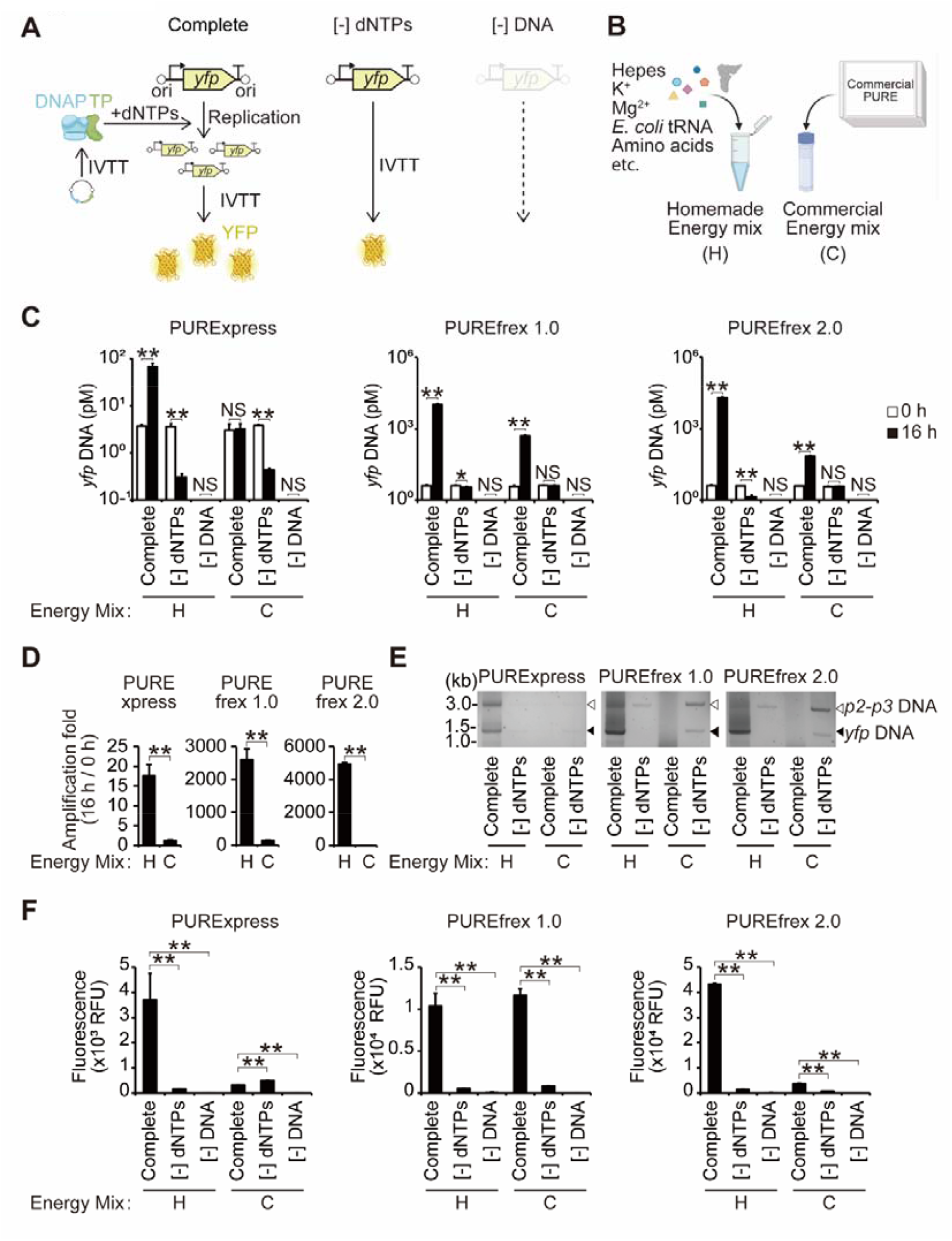
Optimized energy mix restores CADGE across PURE platforms. **(A)** Schematic of CADGE. A linear *yfp* DNA template flanked by *ori* from both sides is amplified by *in situ*-expressed Φ29 DNAP and TP proteins from plasmid DNA via IVTT in PURE. In the presence of dNTPs (Complete), protein-primed DNA replication increases *yfp* template copy number and enhances YFP expression. Without dNTPs ([−] dNTPs), no replication occurs and YFP is expressed only from the input template. Without DNA template ([−] DNA), no YFP expression is detected. **(B)** A schematic of chemically defined homemade mix (H) and commercial mixes (C) supplied with PURExpress or PUREfrex systems. **(C)** Quantification of *yfp* DNA by dPCR at 0 h and 16 h of CADGE reactions. **(D)** DNA amplification fold change (16 h / 0 h) calculated from dPCR measurements in Fig. 1C. **(E)** Estimation of DNA size recovered from CADGE reactions using agarose gel electrophoresis of PCR-amplified DNA. Full-length *ori-yfp* product: 1486 nt. The 3214-nt band DNA is an artifact originating from the plasmid DNA encoding DNAP and TP. **(F)** Estimation of *in vitro* expression of YFP in CADGE reactions by endpoint fluorescence measurements at 16 h. Data are presented as mean + SD (t -test, ** P < 0.01; * P < 0.05; NS, not significant; n = 3). Created in BioRender. Lab, A. (2026) https://BioRender.com/geukhbp.

Quantification of *yfp* DNA by digital PCR (dPCR) before and after the 16-hour incubation of IVTTR reactions at 30°C revealed that reactions containing commercial energy mixes allowed only limited DNA amplification across PURE platforms, with PUREfrex 1.0 showing highest DNA amplification at around two orders of magnitude (Fig. 1C, 1D). In contrast, the optimized, homemade, energy mix markedly enhanced DNA amplification in PUREfrex systems, yielding at least three orders of magnitude increase (Fig. 1C, 1D), while PURExpress enabled modest replication around one order of magnitude (Fig. 1C, 1D). Note that PURExpress showed an approximately 10-fold DNA degradation within 16 hours of incubation regardless of the energy mix used (Fig. 1C), possibly due to previously reported nuclease activity in this system,^43^ although this degradation appears to be lot-dependent in our experience. A smaller, but still significant degradation of *yfp* DNA was observed in PUREfrex 2.0 supplemented with the homemade energy mix (Fig. 1C; ∼3-fold reduction), but not with the commercial energy mix. In contrast, PUREfrex 1.0 showed no comparable decrease.

To verify the integrity of replicated DNA, the *ori-yfp* DNA was amplified by PCR using primers targeting *ori’*s, and the amplified DNA was visualized by agarose gel electrophoresis. Across PURE platforms, the full-length *ori-yfp* amplicon (1.5 kb) was observed in complete CADGE reactions supplemented with the homemade energy mix (Fig. 1E, Complete). Moreover, the amount of recovered DNA from IVTTR reactions with the homemade energy mix was considerably higher than from controls lacking dNTPs (Fig. 1E), thus improving DNA recovery from low concentrations of IVTT template. Unexpectedly, in IVTTR reactions supplemented with the commercial energy mix, PCR amplification yielded higher apparent *yfp* DNA levels in control reactions lacking dNTPs than in complete reactions (Fig. 1E). This counterintuitive result likely reflects reduced PCR efficiency of replicated DNA molecules, as Φ29 DNAP generates products covalently linked to terminal protein (TP) at their 5′ ends, which may sterically hinder the elongating DNA polymerase approaching the 5′ ends of the template. However, the high level of DNA amplification observed in reactions with the homemade energy mix likely overcomes this inhibition. Higher molecular band (∼3.2 kb, Fig. 1E) is an artefact that corresponds to the PCR amplicon from the DNAP and TP-expressing plasmid, which similarly contains Φ29 *ori*’s. The *ori*’s on the plasmid DNA encoding DNAP and TP are not required in CADGE, and should be removed prior to the start of directed evolution, which would eliminate the artefact. Together, these results demonstrate that a chemically defined, replication-optimized energy mix restores and markedly enhances IVTTR performance across PURE platforms.

To validate that DNA replication is compatible with *in situ* cell-free transcription and translation of the amplified gene, we performed endpoint YFP fluorescence measurements. Across platforms, reactions supplemented with the optimized energy mix generally produced higher YFP fluorescence at 16 hours at 30°C than those containing the corresponding commercial energy mixes, except for PUREfrex 1.0, where fluorescence remained comparable to the commercial formulation. In all three systems, YFP fluorescence increased significantly in the presence of both template DNA and dNTPs, confirming that the observed signal reflects DNA replication-coupled transcription-translation (Fig. 1F). Together, these results indicate that while commercial PURE formulations are highly optimized for protein synthesis, the homemade energy mix is more suitable for IVTT-coupled DNA replication in CADGE.

### Optimized energy mix moderately improves protein-primed self-replication

We next tested the performance of the optimized energy mix in self-replication of a minimal genetic module encoding Φ29 DNAP and TP (*ori*-*p2-p3*)^45,52^ (Fig. 2A). In this construct, the *p2* and *p3* genes are encoded on a single linear DNA template flanked by Φ29 *ori*’s, enabling DNAP and TP to drive autocatalytic replication of the DNA encoding them. Since PUREfrex demonstrated significantly higher replication than PURExpress in the CADGE setup, we proceeded to test self-replication only in PUREfrex. DNA replication from an initial template concentration of 2.3 nM was quantified via dPCR after 16 hours of incubation at 30°C in PUREfrex 1.0 and PUREfrex 2.0. In both platforms, moderate amplification of *p2-p3* DNA (10-40-fold) was observed with either energy mix (Fig. 2B). Omission of dNTPs in the IVTTR reactions completely abolished DNA amplification, confirming that the observed increases in DNA concentration were IVTTR-dependent.

**Figure 2.**
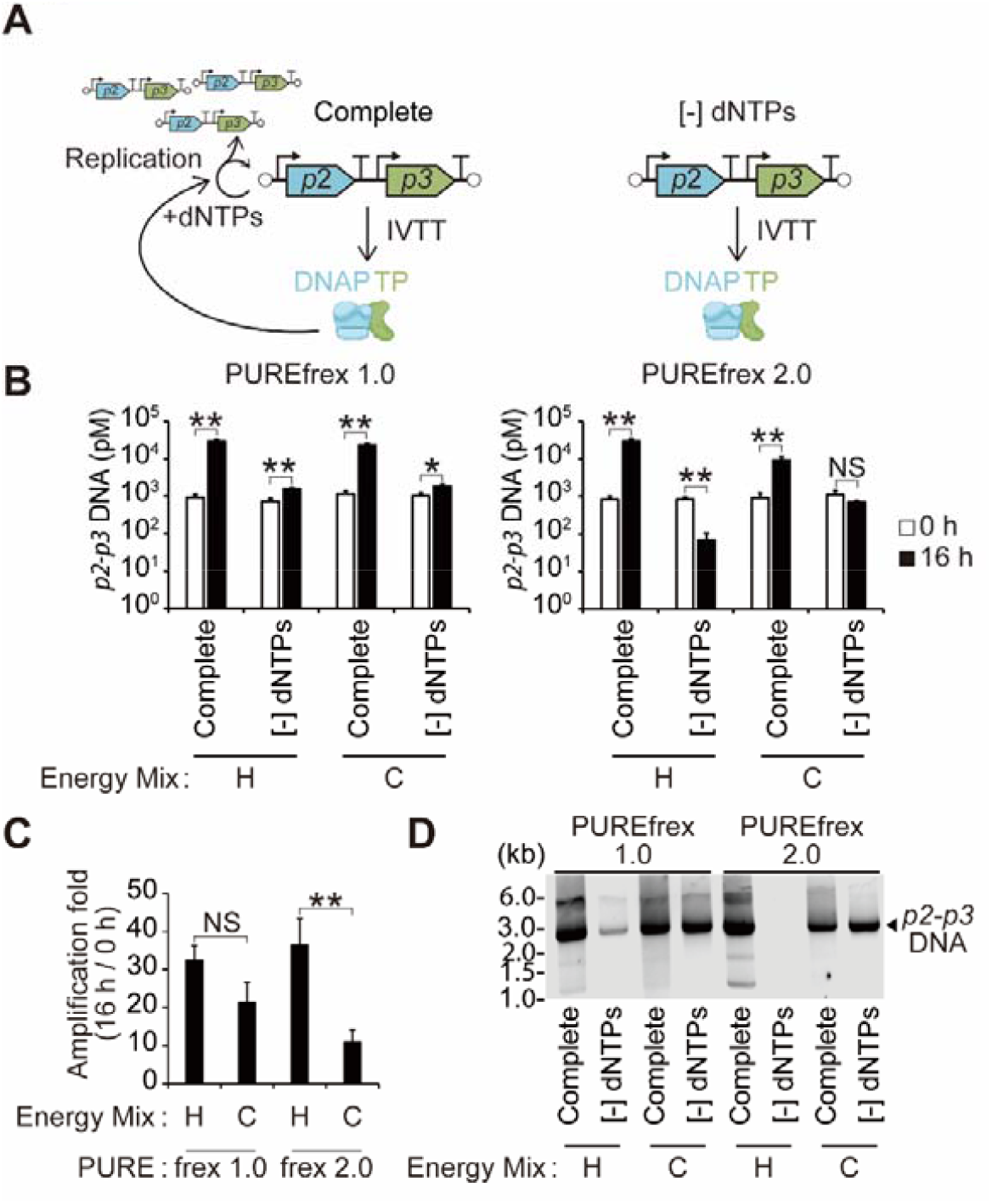
Optimized energy mix supports Φ29 self-replication. **(A)** Schematic of self-replication. *p2* (DNAP) and *p3* (TP) are expressed via IVTT and form an active DNAP–TP complex that initiates replication at flanking *ori* sequences in the presence of dNTPs (Complete), driving autocatalytic amplification. In [−] dNTPs, replication is blocked and only expression from the input template occurs. **(B)** Quantification of *p2-p3* DNA by dPCR at 0 and 16 h in PUREfrex using homemade (H) or commercial (C) energy mixes. **(C)** Amplification fold change (16 h / 0 h) derived from Fig. 2B. **(D)** Agarose gel electrophoresis of PCR-amplified DNA recovered from self-replication reactions. Data are presented as mean + SD (t -test, ** P < 0.01; * P < 0.05; NS, not significant; n = 3). Created in BioRender. Lab, A. (2026) https://BioRender.com/geukhbp.

Comparison of amplification efficiency revealed that the PUREfrex 2.0 supplemented with the homemade energy mix produced higher replication yields than the commercial energy mix, whereas no significant difference was observed in PUREfrex 1.0 (Fig. 2C). This pattern mirrors the weaker enhancement seen for CADGE in PUREfrex 1.0 compared to PUREfrex 2.0 (Fig. 1D), and may reflect the higher translational flux of PUREfrex 2.0. Agarose gel electrophoresis confirmed the accumulation of full-length *p2-p3* DNA products (3.2 kb) in the presence of dNTPs in PUREfrex 2.0 supplemented with the homemade energy mix (Fig. 2D).

These results demonstrate that the PUREfrex 2.0 supplemented with the replication-optimized energy mix enables improved self-replication of a minimal Φ29 DNA replication module compared to the fully commercial formulation. Overall, we show that a DNA replication-optimized energy formulation originally developed for rolling-circle replication can be directly applied to protein-primed Φ29 replication from linear DNA templates. Our study provides reaction conditions that could support selection- and evolution-based experiments and offers a starting point for their further development.

## Methods

### DNA template preparation

Plasmids, primers, and template sequences are listed in Supplementary Tables S1–S3. Linear *ori-yfp* and *ori*-*p2-p3* DNAs were amplified from plasmids G365^28^ and G340^52^ using the same 5′-phosphorylated primer pair (primers 1 and 2). PCR (50 µL) contained 1x Phusion HF buffer, 200 µM each dNTP, 1 µM each primer, 10 ng template DNA, and 0.5 µL Phusion High-Fidelity DNA Polymerase (Thermo Fisher). Cycling conditions were 98°C for 30 s; 25 cycles of 98°C for 10 s, 60°C for 30 s, and 72°C for 15–30 s per kb (1 min 30 s for *yfp*; 3 min for *p2-p3*); followed by 72°C for 10 min. PCR products were purified using the QIAquick PCR & Gel Cleanup Kit with two washes in Buffer PE and elution in 30 µL nuclease-free water.

### Energy mix preparation

Our homemade energy mix was based on PURErep 10x energy formulation^42^, optimized for rolling circle replication in PURE. The 10x mixture contained the following components: 3.6 mM of each of the 20 natural L-amino acids, 700 mM potassium glutamate, 3.75 mM spermidine, 250 mM creatine phosphate potassium salt, 5.18 g/L *E. coli* tRNA, 1 M HEPES–KOH (pH 8.0), 79 mM hemi-magnesium glutamate, and 60 mM dithiothreitol. tRNA was purified from *E. coli* strain A19 following the procedure described ipreviously.^54,55^

### CADGE reaction

The CADGE reaction was adapted using conditions previously optimized for rolling-circle replication.^43^ All reactions were performed in 10 µL volumes and incubated at 30°C for 16 hours in PCR tubes using a thermal cycler, followed by quenching on ice.

Reactions using PUREfrex 2.0 (GeneFrontier) supplemented with homemade energy mix consisted of 1 µL 10x homemade energy mix, 0.5 µL solution I, 0.5 µL solution II, 0.5 µL solution III, 18.75 mM ATP, 12.5 mM GTP, 6.25 mM UTP, 6.25 mM CTP, 20 mM ammonium sulfate, 300 µM dNTPs, 375 µg/mL purified Φ29 SSB, 105 µg/mL purified Φ29 DSB, and 0.6 U/µL SUPERase·In RNase inhibitor (Thermo Fisher), along with 1 nM *p2*-*p3* plasmid (G340) and 10 pM *ori-yfp* linear reporter DNA. Notably, Commercial energy mix (i.e. solution I) was added at 0.1x, primarily to supply the formyl donor (10-formyl-5,6,7,8-tetrahydrofolic acid (THF)), which is required for Met-tRNA^fMet^ formylation.^43^

CADGE using the PUREfrex 2.0 with the commercial energy mix differed from the above reaction conditions in that no 10x homemade energy mix was added, and the reaction contained 5 µL solution I, while keeping the rest of the reagents as mentioned above.`Click or tap here to enter text`.

In reactions using PUREfrex 1.0, the components corresponding to PUREfrex 2.0 solutions I–III were substituted with their respective counterparts,^43^ except that 0.25 µL of PUREfrex 1.0 solution III was added, maintaining the same proportional adjustment as in the PUREfrex 2.0 reactions. For PURExpress (NEB), solutions A (energy, 2.5x stock, final 0.1x) and B (enzymes and ribosomes, 3.33x stock, final 1x) replaced PUREfrex solutions I– III.^42,43^

### Fluorescence quantification

YFP fluorescence (9 µL IVTTR reaction) was measured in 384-well black plates on a BioTek Synergy H1 at 30°C (Ex 513 nm; Em 550 nm to minimize excitation-emission overlap).

### DNA quantification by dPCR

The IVTTR reaction mixtures were diluted 100-fold in TE buffer (10 mM Tris-HCl, 1 mM EDTA, pH 8.0) and stored at −20°C until use. These samples were further diluted in nuclease-free water so that the expected target occupancy (λ) was between 0.6 and 1.6 copies per microchamber, which keeps a sufficient fraction of partitions negative for accurate calculation of DNA concentration. Target-specific primer-probe sets for *yfp* and *p2-p3* were custom-synthesized as FAM-labeled TaqMan Gene Expression Assays (Thermo Fisher) and dPCR was performed using QuantStudio Absolute Q Digital PCR System (Thermo Fisher).

### IVTTR product size estimation by agarose gel electrophoresis

To estimate the size of *yfp* or *p2-p3* DNA amplified in IVTTR reactions, 1 µL of 100-fold diluted reaction mixture was subjected to PCR (100 µL) under the same conditions as for DNA template preparation. PCR products were loaded onto 1% agarose gel without purification and visualized under UV illumination after ethidium bromide staining.

### Self-amplification reaction

Self-replication was conducted under the same bulk CADGE conditions, but without *yfp* reporter DNA and using 2.3 nM *ori*-*p2-p3* linear DNA (pre-amplified by PCR with phosphorylated primers) instead of G340 plasmid.

### Statistical analysis

All experiments were performed in three independent replicates. Data are presented as mean ± SD. Statistical significance was determined by unpaired two-tailed Student’s t-test (P < 0.01 (**); P < 0.05 (*); not significant (NS)).

## Author Contributions

Z.A. conceptualized the research and acquired funding. R.N. performed the experiments.

R.N. and Z.A. designed the experiments. R.N., C.C.R., and Z.A. analyzed the data and wrote the manuscript.

## Acknowledgments

We thank members of the Abil laboratory for helpful discussions and feedback on the manuscript. We are grateful to Miguel de Vega (Centro de Biología Molecular Severo Ochoa, Madrid) for kindly providing purified SSB and DSB proteins. We thank Christophe Danelon for the kind sharing of the G365 and G340 plasmids and for helpful discussions. This work was supported by startup funding from the University of Florida College of Liberal Arts and Sciences and the University of Florida Gatorade Award. C.C.R was supported by the University of Florida Biology Department’s 2025 Graduate Student Opportunity Award. Images were created in BioRender Lab, A. (2026) https://BioRender.com/geukhbp.

