## Supplementary Tables S1-S3 for "CADGE 2.0, Transcription-Translation-Coupled DNA Replication is Improved in a Chemically Modified Cell-Free System"

**Supplementary Table S1. Plasmids used in this paper**

| # | Name | Genetic information | Ref. |
| --- | --- | --- | --- |
| 1 | G340: p2-p3 | oriL191-T7 promoter-SD-p2-VSV terminators-T7 promoter-SD-p3-T7 terminator-oriR194, AmpR | ^1^ |
| 2 | ZA001 (G365): yfp | oriL191-T7 promoter-SD-yfp-T7 terminator-oriR194, AmpR | ^2^ |

**Supplementary Table S2. Primers used in this paper**

| # | Name | Sequence | Description |
| --- | --- | --- | --- |
| 1 | 009ZA | AAAGTAAGCCCCCACCCTCACATG | Forward primer for OriL191 (5' end) |
| 2 | 010ZA | AAAGTAGGGTACAGCGACAACATACAC | Reverse primer for OriR194 (3' end) |

**Supplementary Table S3. Sequence of linear DNA templates for IVTTR**

| *Ori*-*yfp* | AAAGTAAGCCCCCACCCTCACATGATACCATTCTCCTAATATCGACATAATCCGTCGATCCTCGGCATACCATGATCAGGGAGGGAAACTACTACTTAATATATCAATCTATAGACCTACTAGATAGGTTTGTCAATGAACAACATAAAACGACACAGAATCCCACGTTTTAGCGCTTCGTCTGTGTCGCATGTgaaatTAATACGACTCACTATA*gggagaccacaacggtttccctctagaaataattttgtttaactttaag*AAGGAGatatacatATGCGGGGTTCTCATCATCATCATCATCATGGTATGGCTAGCATGACTGGTGGACAGCAAATGGGTCGGGATCTGTACGACGATGACGATAAGGATCCG**ATGGTTAGCAAAGGCGAAGAACTGTTTACGGGCGTGGTGCCGATTCTGGTGGAACTGGACGGCGACGTGAACGGTCACAAATTCAGCGTTTCGGGCGAAGGTGAAGGCGATGCGACCTATGGTAAACTGACGCTGAAATTTATTTGCACCACCGGTAAACTGCCGGTGCCGTGGCCGACCCTGGTTACCACGTTTGGTTATGGCCTGCAGTGTTTCGCGCGCTACCCGGATCATATGAAACAACACGACTTTTTCAAATCTGCCATGCCGGAAGGTTATGTGCAGGAACGTACGATTTTCTTTAAAGATGACGGCAACTACAAAACCCGCGCAGAAGTCAAATTTGAAGGTGATACGCTGGTGAACCGTATTGAACTGAAAGGCATCGATTTCAAAGAAGACGGTAATATCCTGGGCCATAAACTGGAATACAACTACAACTCCCACAACGTTTACATCATGGCAGATAAACAGAAAAACGGTATCAAAGTCAACTTCAAAATCCGCCATAACATCGAAGATGGCTCAGTGCAACTGGCTGACCACTACCAGCAAAACACCCCGATCGGTGATGGCCCGGTTCTGCTGCCGGACAATCATTATCTGAGCTACCAGTCTGCACTGAGTAAAGATCCGAACGAAAAACGTGACCACATGGTCCTGCTGGAATTTGTGACGGCGGCTGGTATTACGCTGGGCATGGATGAACTGTATAAATGA**aagcttcccgggaaagtatatatgagtaaagatatcgacgcaactgaatgaaatggtgaaggacgggtccaggtgtggctgcttcggcagtgcagcttgttgagtagagtgtgagctccgtaactagtcgcgtcgatatccccgggCTAGCATAACCCCTTGGGGCCTCTAAACGGGTCTTGAGGGGTTTTTTG*CCTCCTATGATTGGTTGTCTTATTACCTTACTTCTATTATAGTATAACATGTTAAACGATAGTTTGTCTACCCTTTTCGACAAATTGATGATAATAAATAGTATAGGTATATAGTCGTGATTTAGTTGTTAGATTCTTGTCGAAGATAGTCGGTCAATGGGGAAATGGTGTATGTTGTCGCTGTACCCTACTTT*  Uppercase letters: oriL191  Single underline: T7 promoter (Boxed G: transcription start site)  Lowercase italic letters: T7 phage gene 10 untranslated leader sequence^*1^  Gray-shaded: Shine–Dalgarno (SD) sequence (i.e. RBS: ribosome binding site)  Double underline: 6xHis tag-T7 tag- Xpress tag  Bold: YFP  Thick underline: T7 terminator  Uppercase italic letters: oriR194  Small letters: additional linker sequences |
| --- | --- |
| *Ori*-*p2*-*p3* | AAAGTAAGCCCCCACCCTCACATGATACCATTCTCCTAATATCGACATAATCCGTCGATCCTCGGCATACCATGATCAGGGAGGGAAACTACTACTTAATATATCAATCTATAGACCTACTAGATAGGTTTGTCAATGAACAACATAAAACGACACAGAATCCCACGTTTTAGCGCTTCGTCTGTGTCGCATGTgaaatTAATACGACTCACTATA*gggagaccacaacggtttccctctagaaataattttgtttaactttaag*AAGGAGatatacat**ATGCCGCGTAAAATGTACAGCTGCGATTTTGAAACGACGACGAAAGTTGAAGATTGCCGTGTCTGGGCCTATGGTTATATGAACATCGAAGACCATTCAGAATATAAAATTGGCAACTCGCTGGATGAATTTATGGCGTGGGTGCTGAAAGTTCAGGCCGACCTGTACTTCCACAATCTGAAATTTGATGGTGCGTTCATTATCAACTGGCTGGAACGTAATGGCTTTAAATGGAGCGCCGATGGTCTGCCGAACACCTATAATACGATTATCTCTCGTATGGGCCAATGGTATATGATTGATATCTGCCTGGGCTACAAAGGTAAACGCAAAATTCATACCGTGATCTATGACAGCCTGAAAAAACTGCCGTTTCCGGTGAAGAAAATTGCGAAAGATTTCAAACTGACCGTCCTGAAAGGCGATATTGACTATCACAAAGAACGTCCGGTTGGTTACAAAATCACGCCGGAAGAATATGCGTACATTAAAAACGATATCCAGATTATCGCAGAAGCTCTGCTGATTCAGTTTAAACAAGGCCTGGATCGCATGACCGCCGGCAGTGACTCCCTGAAAGGTTTCAAAGATATCATCACCACGAAAAAATTTAAGAAAGTGTTCCCGACCCTGAGCCTGGGTCTGGATAAAGAAGTTCGTTATGCATACCGCGGCGGTTTTACGTGGCTGAACGACCGTTTCAAAGAAAAAGAAATTGGCGAGGGTATGGTCTTTGATGTGAATAGTCTGTATCCGGCTCAGATGTACTCCCGCCTGCTGCCGTATGGCGAACCGATCGTTTTCGAGGGTAAATATGTCTGGGATGAAGACTACCCGCTGCATATTCAGCACATCCGTTGTGAATTTGAACTGAAAGAAGGCTATATTCCGACCATTCAAATCAAACGTAGCCGCTTCTATAAGGGTAACGAATACCTGAAAAGCTCTGGCGGTGAAATCGCAGACCTGTGGCTGAGTAACGTCGATCTGGAACTGATGAAAGAACATTACGATCTGTACAACGTTGAATACATCTCCGGCCTGAAATTTAAAGCCACCACGGGTCTGTTTAAAGATTTCATTGACAAATGGACCTACATCAAAACCACGTCTGAAGGTGCAATCAAACAGCTGGCTAAACTGATGCTGAACAGCCTGTATGGCAAATTTGCATCTAATCCGGATGTTACCGGTAAAGTCCCGTACCTGAAAGAAAATGGCGCTCTGGGTTTTCGCCTGGGCGAAGAAGAAACCAAAGATCCGGTGTATACGCCGATGGGTGTTTTCATTACCGCGTGGGCACGTTACACCACCATCACCGCCGCACAAGCGTGCTATGACCGCATTATCTACTGTGATACCGACTCAATTCATCTGACCGGCACGGAAATCCCGGATGTGATTAAAGATATCGTTGACCCGAAAAAACTGGGTTATTGGGCACACGAATCGACCTTTAAACGTGCTAAATACCTGCGCCAGAAAACGTACATCCAAGACATCTACATGAAAGAAGTCGATGGCAAACTGGTGGAAGGTTCACCGGATGACTATACCGACATTAAATTTTCGGTGAAATGCGCCGGCATGACCGATAAAATTAAGAAAGAAGTGACGTTCGAAAATTTCAAAGTGGGTTTCAGTCGCAAAATGAAACCGAAACCGGTCCAAGTTCCGGGCGGCGTTGTGCTGGTCGATGACACCTTCACGATCAAATAA**gaattgtactagagTATCTGTTAGTTTTTTTCtactagagtactagagTATCTGTTAGTTTTTTTCatcggatcccgggcccgtcgactgcTAATACGACTCACTATA***gggccctctggagacaccagagggtttacatgtttatttgtttaactttaag***AAGGAGatatacta***ATGGCACGCAGCCCGCGCATCCGCATCAAAGATAACGACAAAGCCGAATACGCCCGCCTGGTGAAAAATACGAAAGCTAAAATCGCACGTACCAAGAAAAAATATGGCGTGGATCTGACGGCTGAAATTGACATCCCGGATCTGGACTCATTTGAAACCCGCGCGCAGTTCAATAAATGGAAAGAACAAGCTAGCTCTTTTACGAACCGTGCGAATATGCGCTATCAGTTCGAGAAAAACGCCTACGGTGTGGTTGCATCGAAAGCTAAAATTGCGGAAATCGAACGTAACACCAAAGAAGTTCAACGCCTGGTCGATGAAAAAATCAAAGCCATGAAAGACAAAGAATATTACGCAGGCGGTAAACCGCAGGGCACGATTGAACAACGTATCGCCATGACCTCACCGGCACATGTGACGGGTATCAACCGTCCGCACGATTTTGACTTCAGTAAAGTTCGCAGTTACTCCCGTCTGCGCACCCTGGAAGAATCCATGGAAATGCGCACGGATCCGCAGTATTACGAAAAGAAAATGATTCAGCTGCAACTGAATTTTATCAAAAGCGTCGAAGGCTCATTTAACTCGTTCGATGCGGCCGACGAACTGATTGAAGAACTGAAGAAAATTCCGCCGGATGACTTTTATGAACTGTTCCTGCGTATTAGCGAAATCTCTTTTGAAGAATTCGATTCTGAAGGCAACACCGTCGAAAATGTGGAGGGTAACGTTTACAAAATTCTGTCGTATCTGGAACAATATCGTCGTGGTGATTTTGATCTGTCGCTGAAAGGCTTCTAA***TAGCATAACCCCTTGGGGCCTCTAAACGGGTCTTGAGGGGTTTTTTG*CCTCCTATGATTGGTTGTCTTATTACCTTACTTCTATTATAGTATAACATGTTAAACGATAGTTTGTCTACCCTTTTCGACAAATTGATGATAATAAATAGTATAGGTATATAGTCGTGATTTAGTTGTTAGATTCTTGTCGAAGATAGTCGGTCAATGGGGAAATGGTGTATGTTGTCGCTGTACCCTACTTT*  Uppercase letters: oriL191  Single underline: T7 promoter (Boxed G: transcription start site)  Lowercase italic letters: T7 phage gene 10 untranslated leader sequence^*1^  Gray-shaded: Shine–Dalgarno (SD) sequence (i.e. RBS: ribosome binding site)  Double underline: 6xHis tag-T7 tag- Xpress tag  Bold: p2  Dashed underline: vesicular stomatitis virus (VSV) terminator^*2^  Lowercase bold italic letters: modified T7 phage gene 10 untranslated leader sequence^*3^  Bold italic: p3  Thick underline: T7 terminator  Uppercase italic letters: oriR194  Small letters: additional linker sequences |

*1: This sequence is derived from the 5′ UTR of phage T7 gene 10, which encodes a very highly expressed phage coat protein, and it forms a characteristic stable stem–loop that strongly enhances translation^3^. The 9-nt epsilon motif (TTAACTTTA) within the leader is perfectly complementary to the 3′ end of *E. coli* 16S rRNA and, being distinct from the SD sequence, enhances mRNA binding to the 16S rRNA and improves translation efficiency^4^. The g10 leader is incorporated in several pET vectors to promote high-level protein expression both in vivo and in in vitro cell-free systems^5^.

*2: This sequence is derived from the vesicular stomatitis virus (VSV) terminator and, despite being only 18 bp long, exhibits ~70% termination efficiency in vitro, comparable to the commonly used T7 terminator^6,7^. Following previous studies^1,8^, we employed two copies in tandem to further enhance transcription termination efficiency.

*3: The native T7 g10 leader cannot be used as the leader sequence for p3, because the native leader, when present upstream of both p2 and p3, creates an 83-bp duplicated region that triggers homologous recombination during IVTTR, generating a parasitic 1.4-kb deletion product that disrupts in vitro evolutionary experiments^1^. We therefore designed, in our previous study, a leader sequence that preserves the RNA secondary structure of the native g10 leader but shares no primary-sequence homology, thereby suppressing recombination while maintaining efficient translation.
